# Diverse competitor B cell responses to a germline-targeting priming immunogen in human Ig loci transgenic mice

**DOI:** 10.1101/2024.01.22.575410

**Authors:** Torben Schiffner, Anne Palser, Joseph G. Jardine, Alessia Liguori, Sergey Menis, Sebastian Raemisch, Xiaozhen Hu, Daniel W. Kulp, Xiaoning Wang, Jolene K. Diedrich, Bettina Groschel, Oleksandr Kalyuzhniy, Ryan Tingle, Khoa Le, Erik Georgeson, Yumiko Adachi, Michael Kubitz, Elise Landais, John R. Yates, James C. Paulson, Devin Sok, Paul Kellam, William R. Schief

**Author notes:** Present address: Cambrian BioPharma, Berlin, Germany (Sebastian Raemisch). Present address: Wistar Institute, Philadelphia, PA, 19104, USA.

## Abstract

Broadly neutralizing antibodies (bnAbs) have promise to protect against HIV infection, but induction of bnAbs by immunization is an unsolved vaccine design challenge. Germline-targeting priming immunogens aim to initiate the induction of bnAbs by specifically activating rare bnAb-precursor B cells that can subsequently be matured using suitable heterologous boosting and shepherding immunogens. Several pre-clinical studies, and the IAVI G001 human clinical trial, have demonstrated the ability of a germline-targeting priming immunogen, eOD-GT8 60mer, to induce precursors of the VRC01 class of bnAbs. However, much less is known about B cells induced against other epitopes of the immunogen. Here, we performed unbiased analysis of B cells induced by eOD-GT8 60mers in Intelliselect Transgenic mice (Kymice) that are transgenic for the human Ig loci and produce human-like BCRs. B cells isolated with intact eOD-GT8 60mer nanoparticles showed a large diversity of non-VRC01-class B cells, with 38% unique clonotypes and only 5% of BCRs belonging to public lineages shared among all animals. We found that many competitors recognize epitopes in close proximity to or overlapping with the VRC01 epitope. These results indicate that optimal boosting of VRC01-class bnAb-precursor B cells primed by eOD-GT8 60mer might require a first-boost immunogen that minimizes recognition of competitor B cells, and such competitors isolated from Kymice could serve as valuable reagents for boost development.

## Introduction

Despite years of intensive research, a vaccine against HIV-1 remains elusive. Most vaccine development approaches aim to induce broadly neutralizing antibodies (bnAbs)^1–3^ that have been shown to provide sterilizing immunity in a number of animal studies^4–10^. However, unmutated germline precursors of bnAbs generally fail to bind to most common variants of the HIV-1 envelope glycoprotein Env, thus making the activation of bnAb precursors unlikely^11^. To overcome this obstacle, a strategy called germline targeting (GT) uses specifically engineered Env variants that bind with high affinity to germline precursors of bnAbs to prime the target B cells, followed by sequential boosting with progressively less mutated Env variants to shepherd affinity maturation towards mature bnAbs^12–15^.

A number of candidate germline-targeting primes has been developed^16–21^, and several have been shown to induce their target B cells in knock-in mouse models^18,20,22–26^. One of the most advanced germline-targeting immunogens is eOD-GT8^27^, an engineered outer domain (eOD) immunogen that contains the conserved binding site for the HIV-1 receptor CD4^16,27^. It has been engineered to bind to precursors of VRC01-class antibodies, which recognize the CD4 binding site (CD4bs) by partially mimicking the interaction of the Envelope protein with CD4^28,29^. For immunizations, the immunogen eOD-GT8 is typically multimerized into a 60-subdomain nanoparticle (termed eOD-GT8 60mer) by fusion to the self-assembling protein lumazine synthase, which results in improved trafficking to lymph nodes and stronger antibody responses^30,31^. eOD-GT8 60mer has been shown to bind to and induce precursors of VRC01-class antibodies in multiple mouse models^22,24,25,32–35^. Recently, eOD-GT8 60mer was administered as recombinant protein with adjuvant AS01B in the G001 clinical trial. The study demonstrated efficient priming of VRC01-class BCRs in 97% of vaccine recipients at substantial frequencies in each individual^36^.

Initial validations of germline-targeting immunogens are often conducted in mouse models with high frequencies of the target B cells^17,20,22–24,32^. However, precursors for many bnAbs are very rare^18,27,37^, and priming of rare B cells requires higher affinities than are required to prime abundant B cell populations^26,34,38–40^. One reason for the requirement for higher affinity might be the intensive competition between B cells in the germinal center reaction^41–45^. While several studies have analyzed the induction of the target B cells, and in some cases epitope-specific competitors^36^, by germline-targeting immunogens, much less is known about B cells induced against other epitopes. Binding studies of sera from G001 clinical trial participants and mice immunized with eOD-GT8 60mer have shown strong antibody responses against eOD-GT8 ^22,24,26,36,38^ and against the lumazine synthase (LS) base nanoparticle^36^, indicating that eOD-GT8 60mer induced antibodies against surfaces outside the CD4bs on both eOD-GT8 and lumazine synthase. Detailed analysis of these competing responses might be important for development of boosting immunogens, because antibodies from primary humoral responses have been shown to modulate secondary responses^46,47^. Further evidence supporting the important role of “off-target” B cells comes from studies using a hyperglycosylated variant of eOD-GT8 60mers, which was designed to suppress “off-target” B cells by occlusion of irrelevant surfaces with immuno-silent glycans. The authors found that glycan masking influenced the relative induction of VRC01-class B cells in vivo^25^. This indicates that competitor B cells are important in shaping B cell responses and knowledge about the precise nature of competing B cell responses might help optimization of germline-targeting immunogens and development of boost immunogens.

We therefore set out to characterize competing B cell responses induced by eOD-GT8 60mers in Intelliselect Transgenic mice (Kymice), transgenic for the complete human Ig loci^48^. Most B cell receptors (BCRs) in Kymice have features similar to those in humans^48^, thus making this an attractive model to predict the most common human antibody responses^49^. A notable exception are VRC01-class precursors, which are substantially less frequent in Kymice than in humans^35,49^. While this makes Kymice non-optimal to investigate VRC01-class responses, they nevertheless present an attractive mouse model to investigate competing B cell responses. Here, we characterized the non-VRC01-class B cell responses induced by eOD-GT8 60mers and investigated their potential to interfere with priming and boosting of VRC01-class BCRs.

## Results

### Serology of non-VRC01-class antibodies induced by eOD-GT8 60mers in human Ig loci transgenic mice

We set out to characterize the overall B cell response to eOD-GT8 60mers in Kymice. Sera from nineteen animals obtained 42 days after immunization with 20 µg eOD-GT8 60mer in Ribi adjuvant (Fig. 1A) showed strong eOD-GT8-specific antibody responses in ELISA with a median EC_50_ of 7.8×10^-4^ (Fig. 1B), consistent with previous results^35^. We also measured antibody responses against the underlying lumazine synthase nanoparticle base. All animals developed strong ELISA responses against naked lumazine (Fig. 1C), similar to results observed in the G001 human clinical trial^36^.

**Fig. 1:**
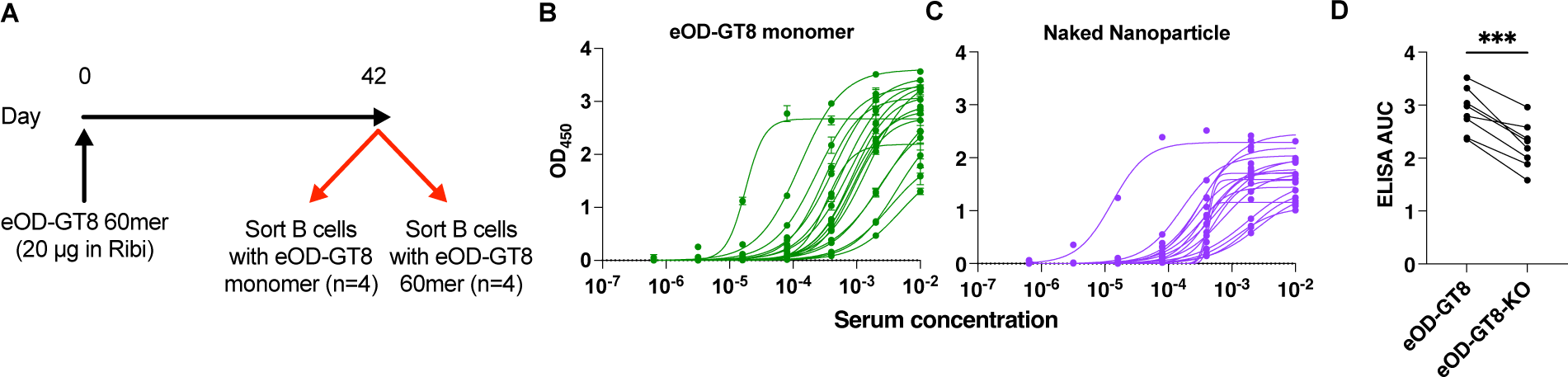
Serum antibody responses of Kymice immunized with eOD-GT8 60mer. **A:** Schematic of the immunization schedule. **B-D**: Total IgG binding of sera from day 42 as determined by ELISA. Raw data of sera from 19 immunized mice biding to eOD-GT8 60mer (**B**), naked lumazine synthase (**C**). **D**: Direct comparison of responses against eOD-GT8 and eOD-GT8-KO. AUC = Area Under the Curve. *** p <0.001 in a paired t test.

We next tested binding of sera from eight animals to eOD-GT8-KO, which contains mutations in the CD4bs that are designed to prevent antibody-binding to this epitope. All animals showed significantly lower binding antibody responses against to eOD-GT8-KO compared to eOD-GT8 (mean 23% reduction in Area under the curve (AUC); p=0.0002 in a paired t test; Fig. 1D), indicating that a substantial proportion of the antibody-responses overlapped at least partially with the CD4bs. Taken together, these data showed that robust IgG responses against both the nanoparticles and the eOD subunit, including the CD4bs, were induced in all animals after a single immunization.

### B cell responses to eOD-GT8 60mers exhibit high intra- and inter-animal diversity

To get a detailed overview of the antibody responses induced by eOD-GT8 60mers, we sacrificed animals 6 weeks after immunization, processed and stained splenic and lymph node cells, and single-cell sorted memory phenotype B cells (GL7-/CD38-IgM-/IgD-; Supp. Fig. 1). Staining included either eOD-GT8 monomer (n=4 animals) or the entire eOD-GT8 60mer nanoparticle (n=4 animals), but by contrast to a previous study^35^, no CD4bs knock-out probe was included. This process enabled us to obtain an unbiased overview of the B cell response to eOD-GT8 60mers. On average, 0.3% and 0.25% of memory B cells bound to the monomer and 60mer, respectively (Fig. 2A). We sequenced BCRs from sorted B cells and obtained sequences of 1173 total HC/LC pairs. eOD-GT8 was specifically engineered to engage VRC01-class B cells, which are commonly defined as antibodies utilizing IGHV1-2 and containing a very short (5-amino acid (aa)) LCDR3^17,25,27,35,37^. We found that 1.4% of BCRs from eOD-GT8-sorted memory B cells utilized IGHV1-2 (Fig 2B). Of these IGHV1-2 sequences, two (11.8%) were paired with a LC containing a 5-aa LCDR3 and were therefore classified as VRC01-class (Fig. 2C). Consequently, in this mouse model, we found that approximately 0.17% of memory B cells induced by eOD-GT8 60mer belonged to the VRC01 class. The remaining IGHV1-2 BCRs were also skewed towards shorter (<9aa) LCDR3 lengths (Fig. 2C). Notably, 35% of IGHV1-2 BCRs had an LCDR3 length of 8aa, and two sequences used IGLV2-23 or IGLV2-8 and were therefore considered potential precursors of the IOMA class, a different class of CD4bs-specific bnAbs^50^.

**Fig. 2:**
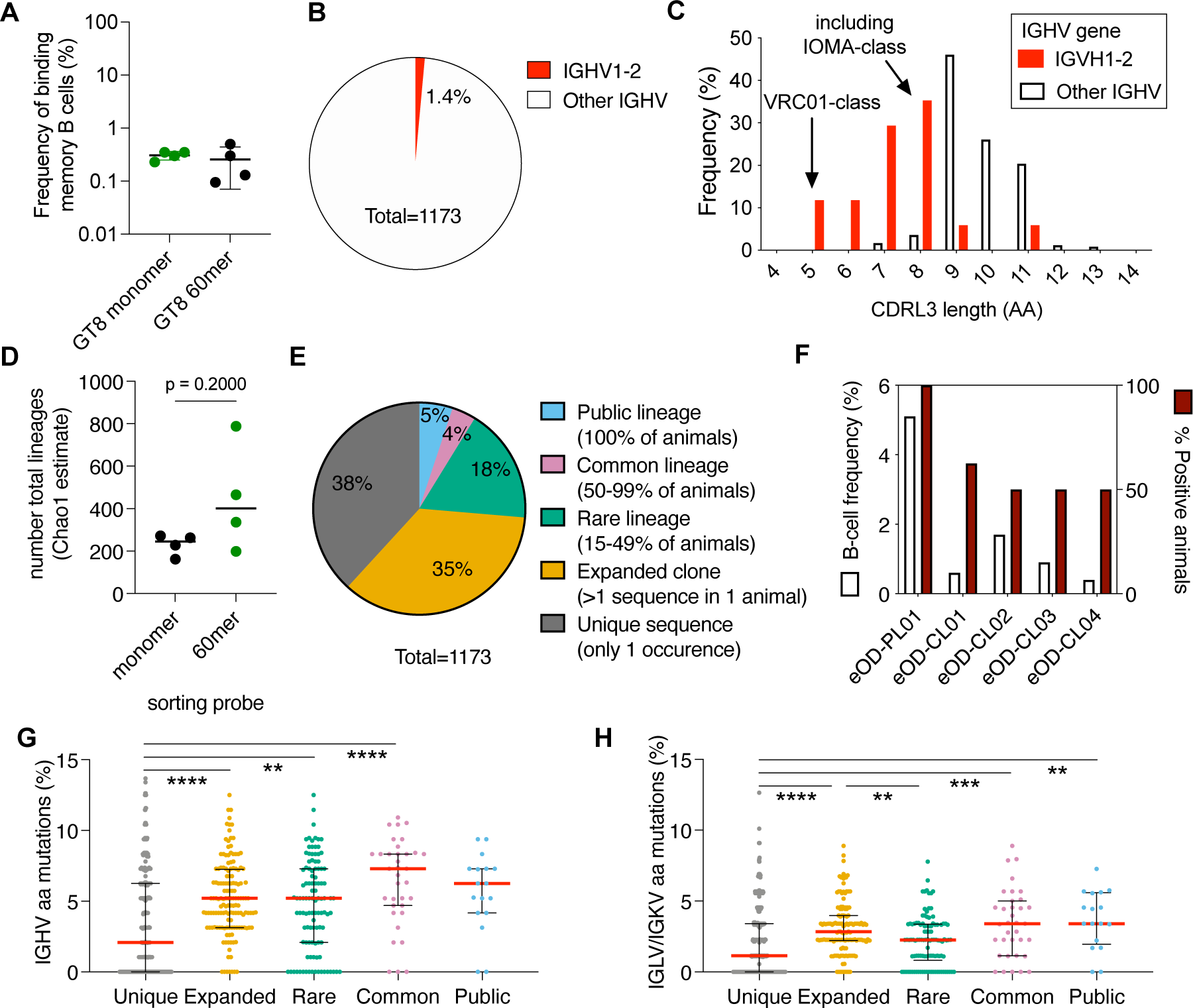
Analysis of BCR sequences induced by eOD-GT8 60mer. Kymice were immunized with eOD-GT8 60mer, and day 42 memory-phenotype B cells (GL7-/CD38-IgM-/IgD-) were sorted and their BCRs were sequenced. **A:** Frequency of B cells binding to the respective sorting probe as a fraction of all memory-phenotype B cells. **B:** Frequency of BCRs utilizing IGHV1-2. **C:** Length distribution of the LCDR3 for IGHV1-2-utilizing BCRs compared to all other sequences. **D:** Chao1 estimation of total B cell lineages recognizing the respective sorting probes. **E:** Similarity of BCRs obtained from different animals. Public lineages were defined as those detected in all animals, common lineages as those found in most but not all animals, rare lineages as those found in more than one but less than half the animals, expanded clones as those found in only one animal but in multiple B cells within that animal, and unique sequences as those found only once in a single animal. **F**. Frequency of common and public lineages among all sequences (left y-axis) and among immunized animals (right y-axis). **G+H:** IGHV (**G**) and IGKV or IGLV (**H**) amino acid mutation levels of the different types of lineages as defined in **E**. Each dot represents the median mutation level of all antibodies from a single lineage. Bars indicate median (red) and interquartile range. ** p <0.01, *** p<0.001, **** p<0.0001; Kruskal-Wallis test with Dunn’s multiple comparison correction. P-values >0.05 are not shown.

To get an overview of the commonly elicited BCRs, we next analyzed the BCR sequences for lineages that could be found in more than one animal. We defined lineages as “similar” if they shared the same IGHV, IGHD and IGLV genes and had the IGHD gene in the same reading frame. Using these criteria, we found 616 different lineages in our dataset. We used a Chao1 estimator^51^, to interpolate the total number of lineages that can be induced against eOD-GT8 60mers in Kymice (Fig 2D). Using this approach, we obtained a lower boundary of at least 1827 eOD-GT8 60mer-specific lineages that can be induced in Kymice. We note that sorting with the intact 60mer nanoparticle tended to recover more total lineages within a single animal (median Chao1 estimate 400) than in animals sorted with eOD-GT8 monomer (median 245), but this difference did not reach statistical significance (p=0.2, Mann-Whitney test). Of the sequenced BCRs, 37% were unique clones that were only found once in one animal (Fig. 2E). An additional 35% of sequences belonged to expanded clones (> 1 sequence) that were found in only one animal (Fig. 2E-F). Expanded clones had significantly higher levels of aa mutations in both heavy chain and light chain V-gene regions (median 5.2% and 2.8% aa mutations in IGHV and IGLV/IGKV, respectively) compared to unique clones (Fig. 2G-F; median 2.1% and 1.12% for IGHV and IGLV/IGKV, respectively; p<0.0001 in a Kruskal-Wallis test with Dunn’s post test). Only 26% of sequences belonged to lineages that could be detected in more than one animal. We identified one public lineage, termed eOD-PL01, that could be isolated from all 8 animals and accounted for 5% of all sequences (Fig. 2E-F). eOD-PL01 used IGHV4-4, IGHD3-9 in the second reading frame, and IGKV1D-39. Further analysis of this lineage showed that the HCDR3 length was 12 or 13aa and most (98%) LCDR3s had a length of 9aa. The most common J genes for heavy and light chain were IGHJ4 and IGKJ4, respectively (both 75%). Sequences perfectly matching these additional CDR-length and J-gene criteria were found in 5 out of 8 animals. eOD-PL01 sequences were highly mutated with median IGHV and IGLV/IGKV aa mutations levels of 6.25% and 3.41%, respectively (Fig. 2G-H). The light chain mutation levels were significantly higher than those found in unique sequences (p<0.01; Kruskal-Wallis test; Fig. 2H).

Further analysis of the sequencing data revealed four lineages that were present in most but not all animals. Frequencies of these common lineages (termed eOD-CL) ranged from 0.4% to 1.7% (Fig. 2F). Lineage eOD-CL01, which was found in 6 out of 8 animals, contained the same IGHV and IGKV genes as the public lineage eOD-PL01. However, it used a different IGHD gene (IGHD3-10 in frame 2) and had a different HCDR3 length (19aa compared to 12-13aa in eOD-PL01) and therefore eOD-CL01 has a fundamentally different HCDR3 than eOD-PL01. eOD-CL03 used the same IGHV and IGHD genes as eOD-CL01, but the D gene was in a different reading frame and the HCDR3 was shorter (13aa vs 19aa). Common lineages showed median aa mutation levels of 7.2% and 3.4% in the IGHV and IGKV/IGKV, respectively, which was comparable to mutation rates in the public lineage but significantly higher than those for unique sequences (p<0.0001; Fig. 2F).

Finally, we defined rare lineages as those found in two or three of the eight animals. Rare lineages made up 18% of all sequences and had mutation levels that were significantly higher than unique sequences (Fig 2G-H; IGHV and IGLV/IGKV mutation levels of 5.2% and 2.3%, respectively; p<0.01; Kruskal-Wallis test). Taken together, these data reveal a highly diverse B cell response to eOD-GT8 60mers, with few lineages shared between different animals.

### B cell responses against eOD-GT8 60mer recognize eOD-GT8 with a wide range of affinities

We next set out to further characterize selected non-VRC01-class lineages and their respective epitopes by expressing, purifying, and characterizing antibodies in vitro. We selected a representative set of antibodies including i) 32 random antibodies sorted with eOD-GT8 60mer, ii) four additional antibodies belonging to eOD-PL01 iii) nine additional antibodies from common lineages, iv) four antibodies from lineages that were only found in 60mer-sorted but not monomer-sorted animals, and vi) 16 randomly chosen unique antibodies. We were able to express and purify 61 of these 65 antibodies.

We initially tested binding of all antibodies in a high-sensitivity BLI assay that utilized the high molecular weight and avidity of the eOD-GT8 60mers to detect even weak interactions of the selected antibodies with the immunogen. In this assay, of 61 antibodies that could be expressed, 56 (93%) bound to the immunogen (Fig. 3A). Using the same assay, we tested binding of all 61 antibodies to naked lumazine synthase 60mer, lacking the eOD-GT8 subunit. Despite the high sensitivity assay setup, none of the antibodies tested showed any interaction with naked lumazine synthase (Fig. 3A). We therefore concluded that most memory B cells induced by eOD-GT8 60mers recognize the eOD-GT8 subunit.

**Fig. 3:**
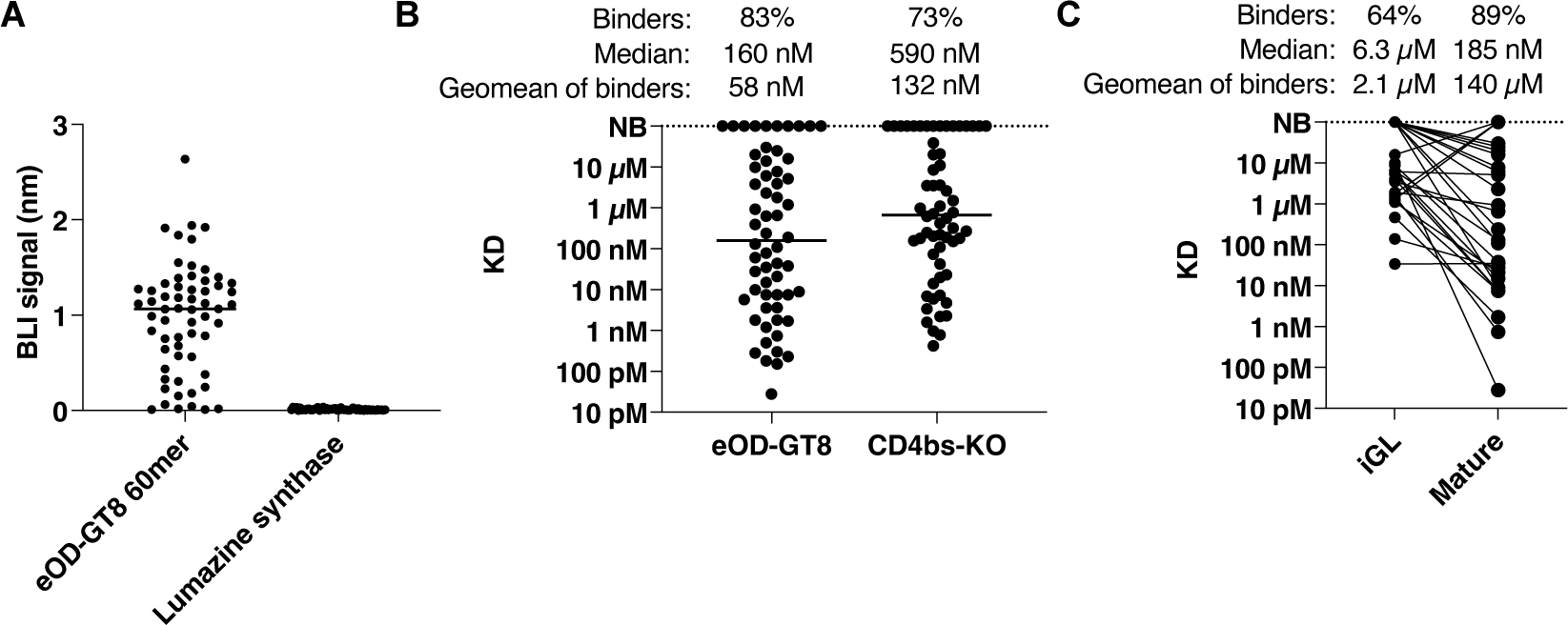
Binding properties of non-VRC01-class antibodies induced by eOD-GT8 60mers. Binding of non-VRC01-class mAbs from Kymice, 6 weeks after immunization with eOD-GT8-60mer, was analyzed by BLI (**A**) and SPR (**B+C**). **A:** Binding of all mAbs to eOD-GT8 and naked lumazine synthase was tested in a high-sensitivity semi-quantitative BLI assay, which utilized the high avidity of the respective nanoparticles to detect even weak binding to the indicated antibodies immobilized on protein A sensor tips. Lines indicate median binding signal. **B**: Monovalent dissociation constants (K_D_s) of all antibodies to eOD-GT8 and eOD-GT8-KO as determined by SPR. Lines indicate median K_D_ values. **C:** Monovalent K_D_s for mature and inferred GL (iGL) antibodies binding to eOD-GT8. NB: No binding.

Next, we performed a kinetic SPR analysis of monomeric eOD-GT8 against all antibodies (Fig 3B). Out of 60 antibodies that were successfully captured onto SPR sensors, 50 (83%) bound with detectable affinity to eOD-GT8, thus confirming the above finding that the majority of recovered BCRs targeted the eOD subunit of the immunogen. Affinities of the binding antibodies were distributed over a wide range, with K_D_ values ranging from 28 pM to 30 µM and with a median K_D_ of 160 nM.

T gain further insights into the affinity maturation of non-VRC01-class antibodies induced by eOD-GT8 60mers, we constructed of inferred-germline (iGL) variants of 35 eOD-GT8 60mer-induced antibodies by reverting the templated regions to their germline sequence. We were able to express and purify 28 pairs of matching mature and iGL antibodies. Of these iGL antibodies, 64% bound to eOD-GT8 with detectable affinity, compared to 89% in the matching mature dataset (Fig. 3C). The geomean affinity of binding antibodies improved from 2.1 µM to 140 nM during affinity maturation. Within the antibody pairs for which reliable K_D_s could be determined, the affinity improved approximately 106-fold during affinity maturation. Taken together, these data showed that B cell responses predominantly targeted the eOD-GT8 subunit and bound with a wide range of affinities.

### Mapping of Epitopes on eOD-GT8

To further map the epitopes of non-VRC01-class B cells induced by eOD-GT8 60mers, we tested binding of mAbs to eOD-GT8-KO, which contains mutations in the CD4 binding site that greatly reduce or eliminate binding by VRC01-class antibodies. SPR analysis showed that 73% of mature antibodies bound to eOD-GT8-KO, with a geomean K_D_ of 132 nM (Fig. 3B). To further characterize the epitopes of these antibodies, we developed a resurfaced variant of eOD-GT8, termed eOD-GT8-RSF. We utilized previous deep mutational scanning data collected during the development of eOD-GT8^27^ to identify surface-exposed mutations outside the VRC01 epitope that showed neutral or positive enrichment upon selection with germline-reverted VRC01-class antibodies. We combined these mutations into a new yeast display library and enriched variants binding with high affinity to germline-reverted VRC01. The best resulting variant, termed eOD-GT8-RSF, had 23 mutations in surface-exposed residues outside of the VRC01 epitope (Fig. 4A). We tested binding of eOD-GT8-RSF to a panel of GL-reverted VRC01-class antibodies and found that binding was no more than 3-fold reduced to any of the antibodies tested, whereas a few mAbs showed considerable (approximately 100-fold) improvement in affinity (Fig. 4B). Therefore, eOD-GT8-RSF had a markedly different surface compared to eOD-GT8 but retained binding to VRC01-class precursors.

**Fig. 4:**
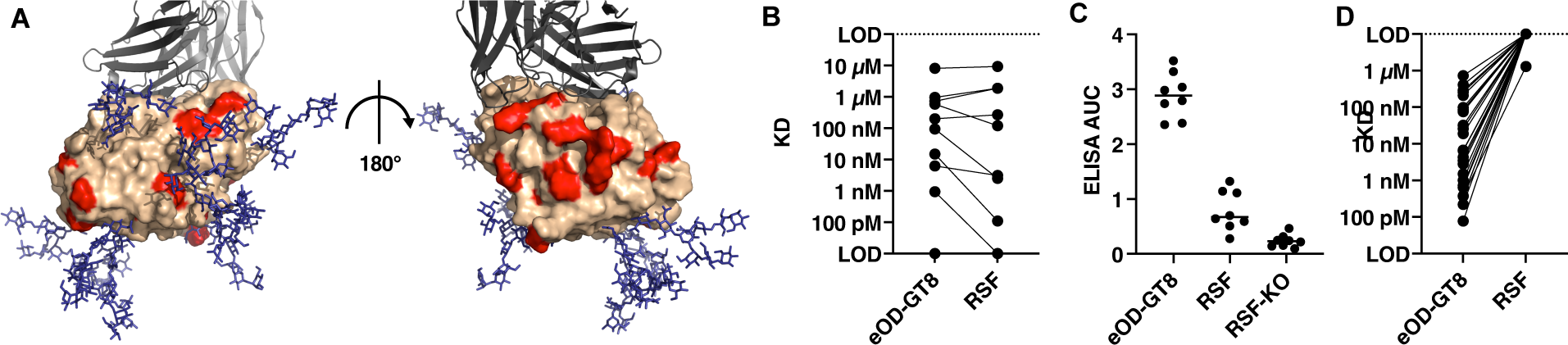
Resurfaced eOD-GT8. **A:** Model of eOD-GT8-RSF (beige) with mutations indicated in red. VRC01-GL is indicated as a grey ribbon model, glycans are shown in blue. **B:** Binding of GL VRC01-class antibodies to eOD-GT8 and eOD-GT8-RSF (RSF) as determined by SPR. Matching antibodies are indicating by solid lines. **C:** ELISA responses from Kymice immunized with eOD-GT8 60mer, showing day 42 serum reactivity to eOD-GT8, eOD-GT8-RSF and eOD-GT8-RSF-KO (RSF-KO). **D:** K_D_s for mAbs from Kymice binding to eOD-GT8 and eOD-GT8-RSF, as determined by SPR. LOD: Limit of detection.

With the resurfaced version of eOD-GT8 in hand, we initially tested binding of sera by ELISA. Binding of week 6 Kymouse sera showed substantially reduced antibody binding to eOD-GT8-RSF (Fig. 4C; 74% reduction on avarage). Introduction of the CD4bs KO mutations into eOD-GT8-RSF further reduced binding of the sera (92% reduction on avarage). We concluded that resurfacing prevented binding of the majority of “off-target” antibodies induced by eOD-GT8 in Kymice.

We next tested binding of the monoclonal non-VRC01-class antibodies induced by eOD-GT8 60mers by SPR (Fig. 4D). As expected, binding of all non-VRC01-class antibodies to eOD-GT8-RSF was completely abolished or severely reduced (>1000-fold increase in K_D_). This included several antibodies that were sensitive to the CD4bs KO mutation, indicating that these antibodies have an epitope that overlaps the CD4bs but is not identical to that of VRC01-class antibodies.

To further map the epitopes targeted by the non-VRC01-class antibodies, we created a panel of patch-revertants of eOD-GT8-RSF, each of which had one surface patch reverted to eOD-GT8. Affinities of GL-VRC01 and VRC01 for all patch-revertants were similar (<4-fold change in KD), confirming that all constructs folded well, and that resurfacing did not have major effects on binding of VRC01-class antibodies (Fig. 5A).

**Fig. 5:**
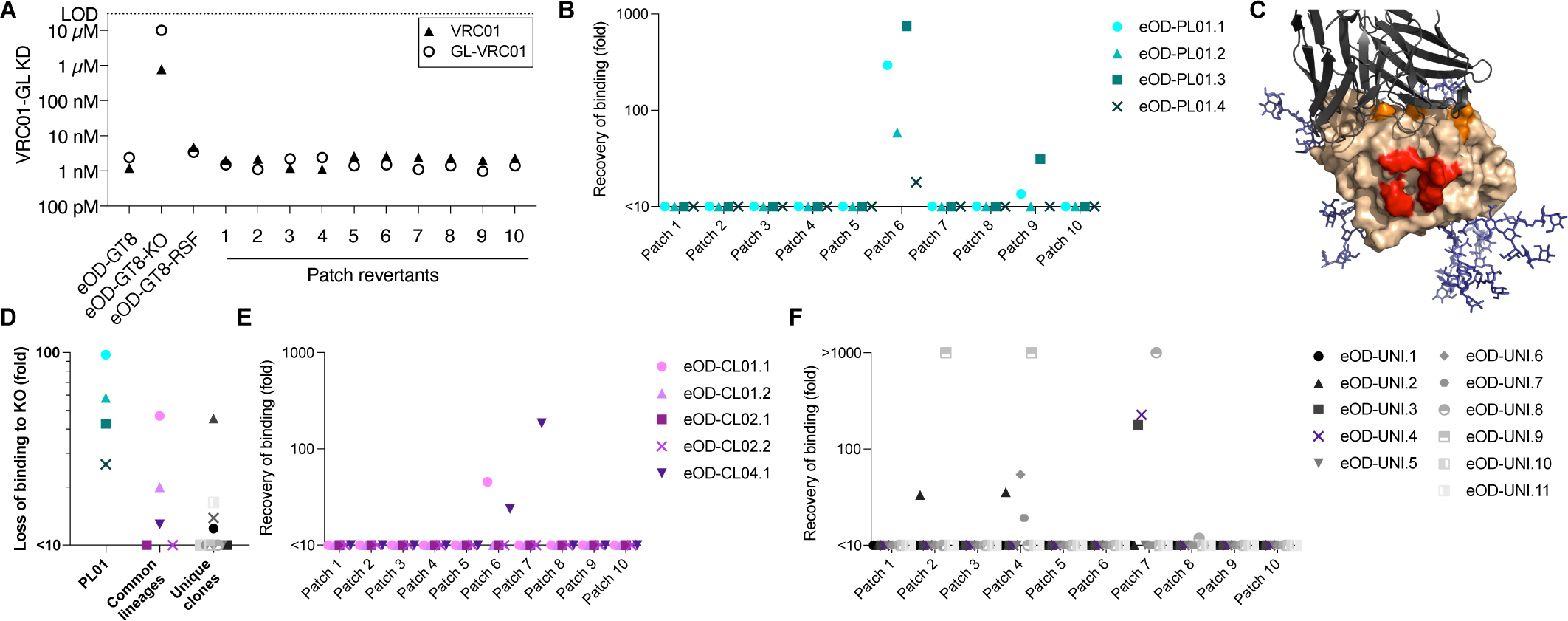
Epitope mapping of non-VRC01-class mAbs induced by eOD-GT8 60mers. **A:** SPR affinities of patch revertants against mature and GL-reverted VRC01. **B:** Mapping of eOD-PL01 class antibodies using the patch revertants. Binding recovery was calculated as the K_D_ for eOD-GT8-RSF divided by the K_D_ for the indicated patch mutant. **C**: Model of eOD-GT8-RSF with patch 6 mutations indicated in red and CD4-KO mutations in orange. VRC01-GL is indicated as a grey ribbon model, glycans are shown in blue. **D**: Loss of binding of eOD-PL01 class antibodies, common lineages and unique clones to CD4-KO to eOD-GT8-KO compared to eOD-GT8. Symbols and colors as in panels **C**, **E** and **F**. **E+F**: Mapping of common lineage antibodies (**E**) and unique clones (**F**) as in **B**.

Using the patch-revertants in combination with the previously described KO mutant, we performed mapping experiments for individual classes of antibodies. For technical reasons, we restricted all further mapping to antibodies that bound strongly (K_D_<1 µM) to eOD-GT8. All tested antibodies from the eOD-PL01 lineage showed negligible binding to resurfaced eOD-GT8-RSF, but binding was largely restored when patch 6 was reverted to eOD-GT8 (Fig. 5B). Patch 6 mutations are located near the stem of loop D (Fig. 5C). All eOD-PL01 antibodies also showed reduced binding to eOD-GT8-KO (median 24-fold increased K_D_; Fig. 5D), indicating that this class recognizes an epitope partially overlapping the CD4bs. Two common lineage antibodies, eOD-CL01.1 and eOD-CL04.1 from common lineages 1 and 4, respectively, were also sensitive to both patch 6 and the CD4bs-KO mutation, indicating that these lineages target an epitope closely related to eOD-PL01 antibodies (Fig. 5D+E). We conclude that eOD-PL01 antibodies and some common lineages target an epitope that overlaps with both the CD4bs and loop D.

Four unique clones could be mapped to patch 4 (Fig. 5F), which centers on loop D (mutations T278M and D279N; HxB2 numbering). Two of these antibodies were also sensitive to mutation T278M alone (patch 2) and insensitive to the CD4-KO mutations, indicating that these antibodies recognize epitopes in close proximity to, but not overlapping with the CD4bs.

Three antibodies (two unique clones and common lineage 4) were mapped to patch 7, which includes mutations in the eOD circular permutation loop (E255V in HxB2 numbering), and sheets β17 (L382F) and β19 (T419R). This surface is largely shielded by the inner domain in gp140 or gp120 Env proteins that include the inner domain. Taken together, these data suggested that most competing B cell responses induced by eOD-GT8 60mer recognized epitopes in loop D and the interface with the inner domain.

### Comparison to the G001 clinical trial

We compared results from the Kymice immunizations with human responses in the G001 clinical trial. However, the sorting in G001 was designed to characterize BCRs specific for the CD4bs, and therefore only lineages sensitive to the CD4-KO mutation were expected to be detected in the G001 dataset. Despite the more restrictive sorting approach, non-VRC01-class lineages were significantly more diverse in the G001 dataset than in Kymice (Fig. 6A). Thirty percent of non-VRC01-class lineages were singlets and a further 31% were expanded clones that were only found in a single individual (Fig. 6B), in line with the results from Kymice. However, no common or public lineages were detected in the G001 dataset. Despite the technical differences in the two studies, these findings provide further support for higher diversity of responses in humans compared to Kymice.

**Fig. 6:**
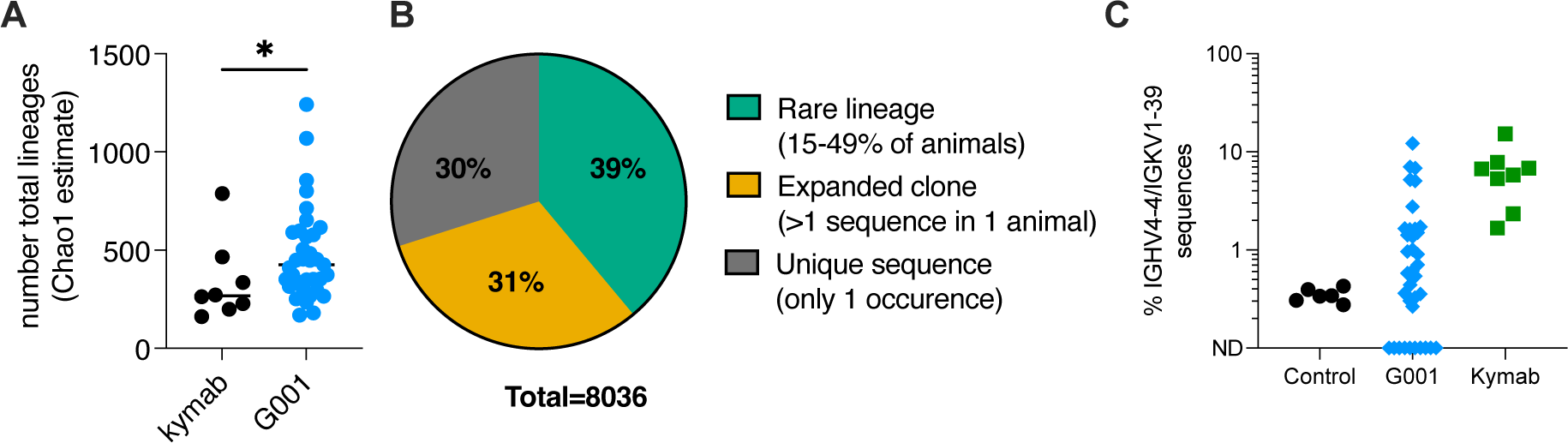
Analysis of non-VRC01-class lineages from the G001 clinical trial. **A:** Chao1 estimation of total B cell lineages detected in the G001 clinical trial upon epitope-specific sorting. **B:** Similarity of BCRs obtained from different donors in G001. Lineage definitions as in Fig. 2E. No public or common lineages were detected. **C**: Frequency of lineages utilizing IGHV4-4 and IGKV1-39 among non-VRC01-class sequences from the G001 clinical trial. Data from available unsorted human control datasets^57,58^ and Kymice are shown for comparison.

As described above, the two most prominent Kymouse lineages that bound with substantially reduced affinity to eOD-GT8-KO, eOD-PL01 and eOD-CL01, both utilized IGHV4-4 and IGKV1-39 but had different HCDR3s (utilizing IGHD3-9 and IGHD3-10, respectively). Sequences using IGHV4-4 and IGKV1-39 were found in 26 donors (74%) and made up 1.3% of non-VRC01-class sequences in G001 (Fig. 6C), indicating a slight enrichment over their estimated baseline frequency (0.34%) that did not reach statistical significance (p=0.3, Mann-Whitney test). The most common IGHD gene among these sequences was IGHD3-3 (43%) rather than IGHD3-9 or IGHD3-10 (both ∼4%) used in eOD-PL01 and eOD-CL01, respectively, indicating that human non-VRC01 class antibodies induced by eOD-GT8 60mer share some but not all features with the lineages discovered in Kymice.

### Hyper-glycosylation of eOD-GT8 abolishes binding of dominant non-VRC01-class lineages

During affinity maturation, competition exists between B cells leading to concerns that the induction of strong non-VRC01-class responses might inhibit the induction of VRC01-class B cells. Duan et al. described a hyperglycosylated version of eOD-GT8^25^, termed eOD-GT8-mut16, and demonstrated that immunization with eOD-GT8-mut16 60mers reduced the frequency of off-target responses in a human V_H_1-2 knock-in mouse model^25^. Based on eOD-GT8-mut16, we developed an improved variant termed eOD-GT8_6G (Fig. 7A), which introduces one additional glycan at position 386 (HxB2 numbering) and uses the amino acid sequence ENET instead of GNGT to introduce a glycan at position 268. Binding of eOD-GT8_6G to VRC01-class iGL antibodies was retained, with none losing more than 3-fold affinity (average 1.6-fold loss) compared to eOD-GT8 (Fig. 7B). Site-specific glycan occupancy analysis confirmed that four of the six new glycosylation sites were predominantly occupied (>50%; Fig. 7C), and one more site was 49% occupied. Of the new sites, only position 380 (HxB2 numbering) was virtually unoccupied (<20% occupancy).

**Fig. 7:**
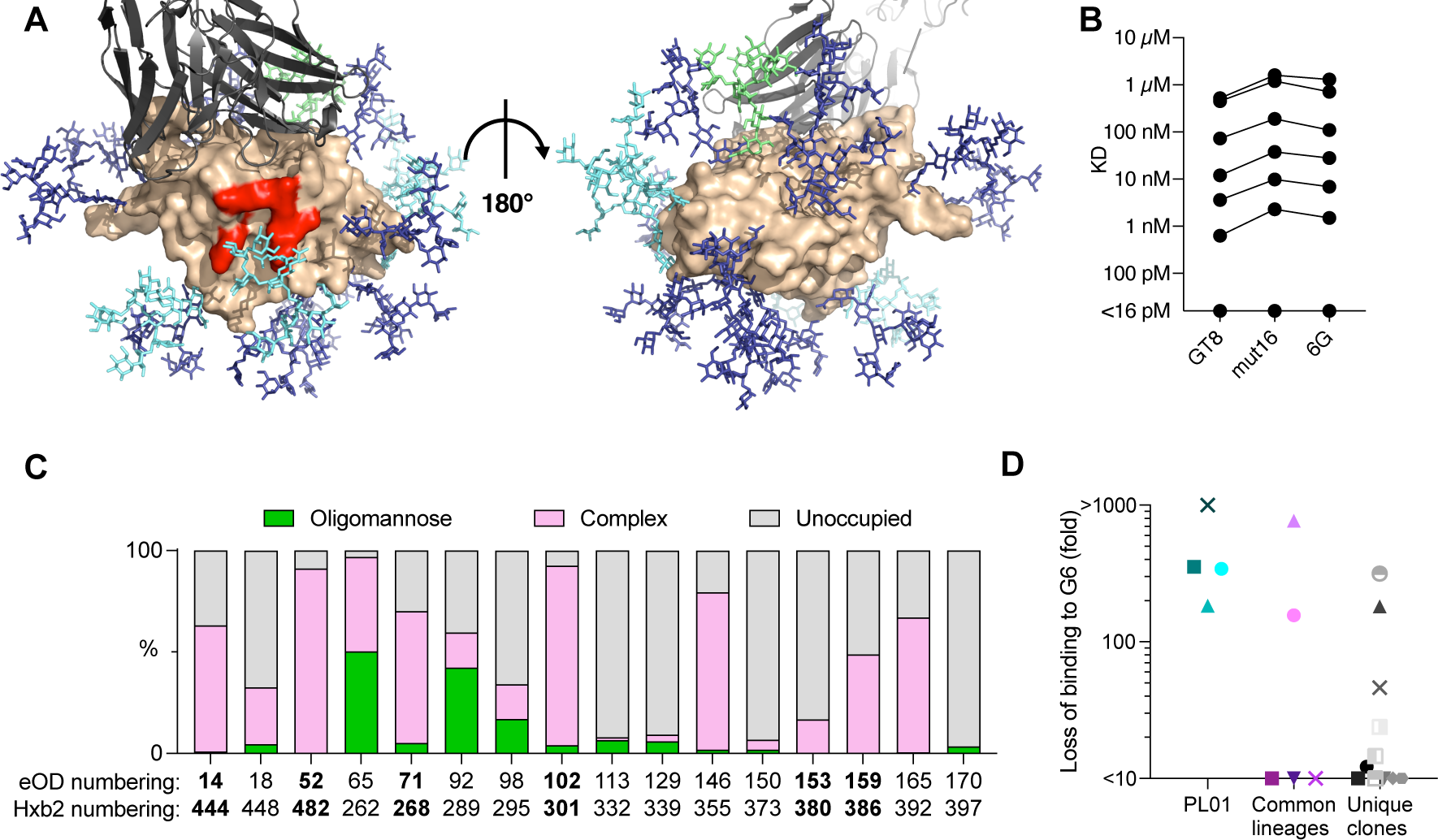
Hyperglycosylation of eOD-GT8. **A:** Model of eOD-GT8-6G (beige) with glycans shown as sticks. Glycans shared with eOD-GT8 are shown in blue, new glycans introduced in eOD-GT8-mut16^25^ are shown in cyan, and the new glycan from 6G in green. VRC01-GL is indicated as a grey ribbon model. **B**: SPR-measured K_D_s for glycan mutants binding to germline-reverted VRC01-class antibodies. **C**: Glycan occupancy as determined by mass spectrometry. Positions are indicated relative to eOD and in HxB2 numbering. New sequons are shown in bold. **D**: Loss of binding of eOD-PL01 class antibodies, common lineages and unique clones to CD4-KO to eOD-GT8-6G compared to eOD-GT8. Symbols and colors as in Fig. 5.

We next tested binding of eOD-GT8_6G to non-VRC01-class antibodies. Among the antibodies that bound tightly (K_D_<1 µM) to eOD-GT8, 60% lost more than 10-fold affinity to eOD-GT8_6G (Fig. 7D). In particular, all eOD-PL01 lineage antibodies lost more than 100-fold affinity. These data show that hyper-glycosylation of eOD-GT8 substantially reduced binding of many non-VRC01-class antibodies and abrogated binding of the dominant competing antibody lineage.

## Discussion

Germline-targeting vaccination strategies aim to induce specific B cell classes, and several vaccine candidates have been proposed that have each been designed to bind with high affinity to precursors for specific bnAb classes^16–20^. However, in pre-clinical experiments, less emphasis has been put on competing “off-target” B cells that have the potential to outcompete the desired lineages during the intense competition of affinity maturation. With few exceptions^35^, most pre-clinical in vivo experiments were conducted in animal models with substantially reduced B cell competition^22–24^, and competing B cells predominantly utilized mouse Ig genes^18,20,22–26^ that differ from the human antibody repertoire. Here, we used the Intelliselect Transgenic mice (Kymice), transgenic for the human Ig loci to investigate global B cell responses to one of the most advanced germline-targeting vaccine candidates, eOD_GT8 60mer, which demonstrated promising results in the G001 human phase 1 clinical trial^36^. It was previously shown that the targeted VRC01-class BCRs are approximately 300 to 900-fold less frequent in this mouse model than in humans^35^, therefore making this model less suitable to test the induction of VRC01-class B cells. In addition, long HCDR3s, which are a requirement for several HIV bnAb classes^52^, have been shown to be less comon in Kymice than in humans^49^, thus complicating testing of bnAb induction in these animals. However, despite differences in the frequencies of specific lineages, the overall BCR repertoire in Kymice mice is comparable to humans^48,49^, therefore making this system suitable to investigate “off-target” B cells.

Previous experiments with eOD-GT8 60mers in this mouse model^35^ or the G001 clinical trial^36^ only characterized CD4bs epitope-specific competitors. Here, we did not use an epitope knock-out probe in our single-cell sorting strategy, thereby gaining an unbiased overview of all memory B cells induced by eOD-GT8 60mer. Sequencing of B cell receptors showed a highly diverse B cell repertoire within each animal. Few of the lineages were shared between different animals, and only one lineage found in every animal. We note that other common lineages might conceivably have been present in every animal but remained undetected in our study due to undersampling. The public and common lineages showed a trend towards higher levels of somatic hypermutation compared to lineages that were detected in only a single or few animals, suggesting that these could have outcompeted other lineages in germinal center reactions, thereby being subjected to more rounds of affinity maturation. Therefore, the public and common lineages might be of particular importance in attempts to modify immunodominance in favor of VRC01-class antibodies.

Previous studies have reported convergent antibody responses upon infection with or vaccination against SARS-CoV-2, but shared antibody lineages made up only small fractions of each individual’s responding BCR repertoire, and each convergent antibody lineage was only found in a minority of individuals, falling into our category of “rare” lineages^53–55^. By contrast to the SARS-CoV-2 spike glycoprotein trimer, which contains 22 glycosylation sites per 140 kDa protomer, the eOD-GT8 60mer is more densely glycosylated, containing 10 glycosylation sites per 37 kDa protomer. Our data showed that the dense glycan shield, which has evolved in response to immune pressure on HIV-1 Env and was retained during development of eOD-GT8, still permitted a large number of B cells to engage eOD. However, despite the diversity of the B cell response, memory B cells still targeted a relatively small number of epitopes. In particular, competing responses were mapped to the CD4 binding site, which has been enlarged in eOD-GT8 by removing the N276 glycan, and de novo surfaces created due to the removal of the inner domain during eOD development^16,27^. The N276 glycosylation site is highly conserved in diverse HIV strains strains^56^, and therefore the underlying surface is shielded from antibody recognition in native Env glycoproteins. Similarly, the de novo surfaces created by removal of the inner domain are occluded on native Env glycoprotein, and have therefore not been subjected to immune pressure from B cell recognition, which might explain their comparatively high immunogenicity when exposed on eOD-GT8 60mers. We note that all antibodies that were successfully mapped in this study recognized surfaces that would be sterically occluded by the inner domain present on previously proposed boosting candidates^32,33^, including native Env trimers and gp120 core immunogens. It is therefore likely that the majority of “off-target” responses induced by eOD-GT8 60mer will not be re-engaged by current candidate booster immunogens.

Approaches to improve the glycan shield have already been shown to reduce competing non-VRC01-class B cell responses in model systems^25^. Here, we showed that the public lineage eOD-PL01, which was detected in every vaccinated Kymouse, binds to an epitope that can be covered with a glycan at position 52 (382 in HcB2 numbering). eOD-PL01 antibodies showed drastically reduced affinities to eOD-GT8-6G, which contains the glycan at position 52. Therefore, the data collected in this study will be helpful in further optimizing the glycan-shield of the eOD-GT8 priming immunogen.

Surprisingly, none of the antibodies identified in this study bound to a naked lumazine synthase construct that represented the base nanoparticle fused with eOD-GT8 to multimerize the immunogen into 60mers. Nevertheless, strong serum antibody responses against naked lumazine synthase demonstrated that such B cells are consistently induced upon immunization. A possible explanation for the discrepancy between the ELISA results and the findings from single B cell sorting might be that B cells targeting lumzine synthase were induced in vivo by partially degraded immunogens that expose lumazine synthase subunits. The sorting probes utilized in this study were largely composed of intact nanoparticles in which the lumazine synthase was sterically occluded by the eOD-GT8 subunits. Further studies that specifically sort with naked lumazine would be required to elucidate the properties of B cells recognizing the underlying nanoparticle.

In summary, we discovered a highly diverse B cell response induced by eOD-GT8 60mers, which was predominantly focused on epitopes in close proximity to or overlapping with the CD4bs. The unbiased characterization of memory B cell responses targeting the entire eOD-GT8 60mer nanoparticle provided complementary data to previous reports restricted to epitope-specific responses and will help guide the development of boosting candidates designed to induce broadly neutralizing antibodies against HIV.

## Acknowledgements

We thank Allan Bradley, Glenn Friedrich, Karen Makar, and Pervin Anklesaria for assistance in arranging the studies. All mice were maintained, and all procedures carried out under United Kingdom Home Office License 70/8718 and with the approval of the Sanger Institute Animal Welfare and Ethical Review Body.

## Funding

This work was supported by: The Bill and Melinda Gates Foundation (BMGF) (to P.K.); Bill and Melinda Gates Foundation Collaboration for AIDS Vaccine Discovery (CAVD) awards INV-007522 and INV-008813 for the IAVI NAC Center (to W.R.S.); National Institute of Allergy and Infectious Diseases (NIAID) awards UM1 AI100663 (Scripps Consortium for HIV/AIDS Vaccine Discovery) and UM1 AI144462 (Scripps Consortium for HIV/AIDS Vaccine Development) (to W.R.S., D.S., and J.C.P.); the Alexander von Humboldt foundation (T.S.); the Helen Hay Whitney Foundation (J.G.J.); the Ragon Institute of MGH, MIT and Harvard (to W.R.S.); and the IAVI NAC (to W.R.S. and D.S.). The research at Kymab, a Sanofi Company was supported by a grant from the Bill and Melinda Gates Foundation, OPP1159947.

## Competing interests

Materials and information concerning the immunogens are available by material transfer agreement from the Scripps Research Institute. IAVI and the Scripps Research Institute have filed a patent (U.S. PCT Application no. PCT/US2016/038162) relating to the eOD-GT8 immunogens in this manuscript, which included inventors J.G.J., D.W.K., S.M., and W.R.S. The Kymab mouse strains described are corporate assets protected by multiple patents; access to these mice is available through licensing.

---

**Supp. Fig. 1:**
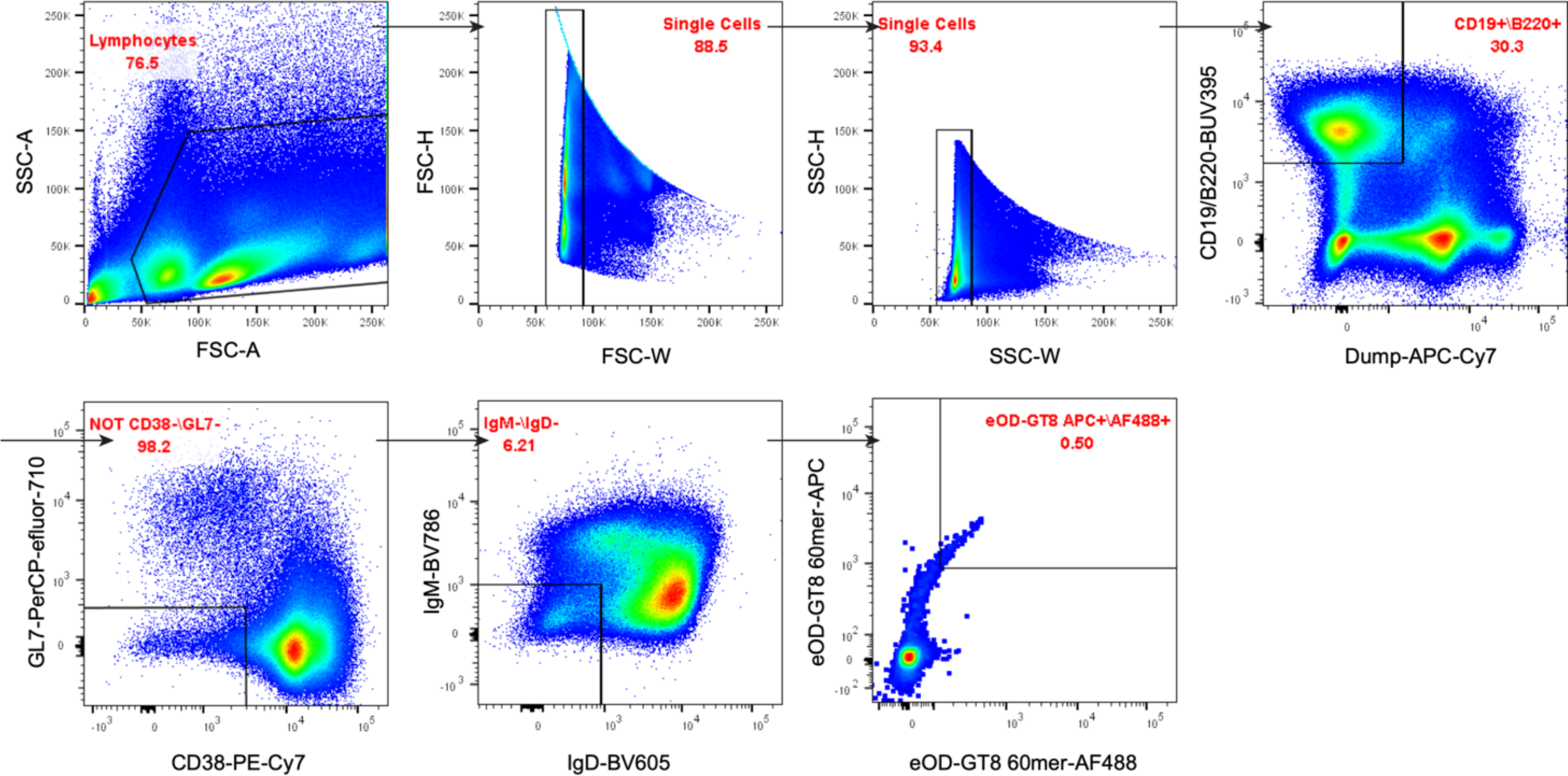
Gating strategy used to sort memory phenotype B cells.

## Methods

### Resurfacing of eOD-GT8

A resurfaced variant of eOD-GT8, termed eOD-GT8-RSF, was developed using directed evolution by yeast surface display as described previously^27^. The yeast library was designed based on deep mutational scanning data used in the design of eOD-GT8^27^. Briefly, mutations of surface-exposed residues that showed a neutral or positive enrichment across the different germline antibody datasets were selected for library incorporation. Emphasis was placed on identifying positions that could accommodate additional N-linked glycosylation motifs. 25 positions were identified, each with 1-3 potential mutations: R8T, S31A, N32R, A49EQ, R50Q, T56N, V58E, E70N, E71N, E72S, V73T, R76T, E78VM, R81M, D82N, K111Q, N114K, Q118D, E125H, G128E, R130K, K135NEQ, E143A, D159N, F169H. A library containing these positions was produced and displayed on the surface of yeast cells. Cells were sorted twice by FACS selecting all clones that bound with comparable affinity to glVRC01 as eOD-GT8. DNA was harvested from the recovered cells and sequenced using next-generation sequencing. Nine enriched variants that contained between 18 and 25 mutations from the original eOD-GT8 were selected for expression and characterization. The final version contained 23 total mutations and introduced two additional N-linked glycosylation sites (N71 and N135). Models of antigens were prepared using AlphaFold^59^, followed by FastRelax^60^ and GlycanTreeModeler^61^ in the Rosetta Macromolecular Modeling software suite^62,63^. Images were generated in Pymol.

### Protein expression and purification

Proteins were expressed and purified as described previously^64^. Briefly, genes were synthesized and cloned into pHLSec or its variant pCWSec by Genscript inc. Proteins were expressed transiently under chemically defined conditions in Freestyle 293F-cells (Thermo Scientific). His-tagged proteins were purified from purified supernatants using immobilized metal affinity chromatography followed by size-exclusion chromatography (SEC). Nanoparticles were purified by *Galanthus nivalis lectin* affinity chromatography (Vector laboratories) followed by SEC. Antibodies were purified by protein A affinity chromatography followed by buffer-exchange into tris-buffered saline.

### Labeling of sorting probes

Avi-tagged proteins were biotinylated using the biotinylated by BirA enzymatic reaction (Avidity, Inc) according to the manufacturer’s protocol and purified by SEC. eOD-GT8 60mer was directly conjugated to fluorochromes using the Alexa Fluor® 488 Antibody Labeling Kit or Alexa Fluor® 647 Antibody Labeling Kit (Thermo Scientific) according to manufacturer’s instructions, except that the buffer contained 20 mM Tris, 4x more protein was used per reaction and the reaction time was reduced to 30 min. Using these optimized conditions resulted in ∼10 fluorophores per 60mer.

### Single-cell sorting and sequencing of immunized Kymice

This study was carried out under Project Licenses 70/8718 issued by the UK Government Home Office under Animal (Scientific Procedures) Act (A(SP)A), 1986, incorporating Directive 2010/63/EU of the European Parliament, and with the approval of the Sanger Institute Animal Welfare and Ethical Review Body. The Institute complied with the Code of Practice issued by the UK Government which aids compliance with the A(SP)A. The Institute has a PHS assurance F16-00128 (WTSI). Intelliselect Transgenic mice (Kymice)^35,49^, (n=8) were immunized with 20 µg eOD-GT8 60mer in Ribi adjuvant (Sigma-Aldrich). 42 days later, animals were sacrificed, and spleens and lymph nodes were processed for single B cell sorting as described previously^27^. Briefly, cells were stained with fluorophore-conjugated antibodies to murine CD4, CD8, F4/80, CD11c, Gr-1, CD19, B220, IgD, IgM, CD38, and GL7 markers. 50 nM of biotinylated sorting probes were coupled to Streptavidin-AF488 and Streptavidin-PE (Life Technologoies) in equimolar ratios, respectively, whereas 60mers were directly conjugated as described above. Memory phenotype B cells were selected for the phenotype CD19+, B220+, CD4-, CD8-, F4/80-, CD11c-, Gr-1-,IgM-, IgD-. B cells of interest were single-cell sorted into 96-well plates containing lysis buffer on a BD FACSAria Fusion sorter and stored at −80°C. Reverse transcription and subsequent PCR amplification of heavy and light chain variable genes were performed using SuperScript III (Life Technologies) as described previously^27^. PCR reactions were performed in 25 μl volume with 2.5 μl of cDNA transcript using HotStar Taq DNA polymerase master mix (Qiagen) and previously described primer mixes. Second round nested-PCR reactions were performed using Phusion proof reading polymerase (NEB). A third round of PCR was performed with HotStar Taq DNA polymerase master mix (Qiagen) using primers with barcodes specific to the plate number and well location as well as adapters appropriate for sequencing on an Illumina MiSeq.

Amplified IgG heavy- and light-chain variable regions were sequenced on an Illumina MiSeq (600-base v3 kit; Illumina) and reads corresponding to the same plate/well location were combined into consensus sequences. Germline assignment and sequence annotation was performed in Abstar (https://github.com/brineylab/abstar).

### ELISA

Enzyme-linked immunosorbent assays were performed by directly coating antigens onto high-binding ELISA (Corning) plates at 2 µg/mL in PBS overnight. Wells were washed with washing buffer (PBS + 0.2% Tween20), and blocked with 1% fetal bovine serum (FBS) + 5% skim milk in washing buffer for 1 h. Serially diluted sera in 1% FBS + washing buffer were added for 1h, plates were washed and anti-mouse IgG secondary antibody diluted 1:2000 in 1% FBS + washing buffer was added for 1h. After washing, wells were developed with TMB-Ultra (Thermo Fisher), reactions were stopped with H_2_SO_4_ and absorption was measured at 450 nm. Data were background-subtracted, and concentrations were log-transformed. EC_50_ and area under the curve (AUC) values were calculated using Graphpad Prism 9.5.1.

### Biolayer interferometry

BLI measurements were performed on an Octet Red96 instrument at 30°C in SPR buffer (HBS-EP+ pH 7.4 running buffer (20x stock from Teknova, Cat. No H8022) supplemented with BSA at 1mg/ml). Protein A sensors were loaded with indicated antibodies for 180 seconds at 5 µg/mL and unbound antibody was washed off with SPR buffer for 30 seconds. Nanoparticles were passed over the sensors for 180 seconds and binding signal was defined as the difference between the signal obtained at the end of the association period and the signal at the baseline between loading and association. Sensors were regenerated using 1.7% phosphoric acid.

### Surface Plasmon Resonance

SPR experiments were performed on a determined on a ProteOn XPR36 (Bio-Rad) instrument using SPR buffer. Approximately 600 RUs of anti-human IgG (Cytiva) were immobilized on a GLC Sensor Chip (Bio-Rad) using amine-coupling. Regeneration solution was 3M magnesium chloride injected four times for 180 seconds per cycle. Antibodies was immobilized for 60 seconds at 1 µg/mL. Dilution series of monovalent analytes were passed over the immobilized antibodies for 120 seconds, followed by a dissociation period of 300 seconds. Raw sensograms were analyzed using interspot and column double referencing in the ProteOn Manager software (Bio-Rad). KDs were calculated using equilibrium fits or kinetic fits with Langmuir model, or both, when applicable.

### Site-specific glycan analysis

Glycan occupancy was analyzed as described previously^65,66^. Briefly, ∼50 µg protein was denatured with 8M Urea in 100 mM ammonium acetate, reduced with 10mM dithiotreihtol at 56°C for 1h and treated with iodoacetamide. After buffer-exchange, proteins were digested with proteases chymotrypsin, trypsin and Arg-C, alone or in combination. Digested proteins were de-glycosylated with EndoH, followed by treatment with PNGase in O^18^-H2O (97%, Sigma-Aldrich). Following separation on 30 cm, 75 µm ID column packed with BEH 1.7 μm C18 resin (Waters), samples were analyzed on a Fusion Orbitrap tribrid mass spectrometer (Thermo Scientific). MS and MS/MS data were extracted from RAW files by using RawConverter^67^ and processed with software package Integrated Proteomics Pipeline-IP2 (version 4)^68,69^ as described previously^65,66^. Each peak was smoothed and fit to Gaussian distribution to calculate the abundance of peptide using peak area. Unidentified peptides were retrieved by retention time, isotope matching scoring, and accurate mass checking.

### Statistical analyses

Statistical analysis using the indicated tests and plotting of all data was performed using Graphpad Prism 9.5.1.

